# Cadherin switch marks germ layer formation in the diploblastic sea anemone *Nematostella vectensis*

**DOI:** 10.1101/488270

**Authors:** E.A. Pukhlyakova, A. Kirillova, Y.A. Kraus, U. Technau

## Abstract

Morphogenesis is a shape-building process during development of multicellular organisms. During this process the establishment and modulation of cell-cell contacts play an important role. Cadherins, the major cell adhesion molecules, form adherens junctions connecting ephithelial cells. Numerous studies in Bilateria have shown that cadherins are associated with the regulation of cell differentiation, cell shape changes, cell migration and tissue morphogenesis. To date, the role of Cadherins in non-bilaterians is unknown. Here, we study the expression and the function of two paralogous classical cadherins, cadherin1 and cadherin3, in the diploblastic animal, the sea anemone *Nematostella vectensis*. We show that a cadherin switch is accompanying the formation of germ layers. Using specific antibodies, we show that both cadherins are localized to adherens junctions at apical and basal positions in ectoderm and endoderm. During gastrulation, partial EMT of endodermal cells is marked by a step-wise downregulation of cadherin3 and upregulation of cadherin1. Knockdown experiments show that both cadherins are required for maintenance of tissue integrity and tissue morphogenesis. This demonstrates that cnidarians convergently use cadherins to differentially control morphogenetic events during development.

## Introduction

Morphogenesis is a process of tissue and organ formation during organism development. (Gilbert, 2013). Morphogenesis is driven by coordinated cell shape changes, cell migration, cell proliferation and cell death and cell adhesion. The key morphogenetic events during early development are gastrulation and germ layer formation, folding of the neural tube and body axis elongation. Cadherins are transmembrane cell adhesion molecules, which play an important role in these processes. They not only provide the mechanical connection between cells, but also control cell-cell recognition, cell sorting, tissue boundary formation, signal transduction, formation of cell and tissue polarity, cell migration, cell proliferation and cell death (Gumbiner, 2005; Halbleib and Nelson, 2006). In adult tissues, cadherins preserve stable and ordered tissue integrity (Angst et al., 2001; Halbleib and Nelson, 2006).

Classical cadherins are conserved molecules present in all animals whose genomes have been analyzed (Alberts, 2007). They are major components of the adherens junctions between cells, which are conserved structures of epithelial cells in most animals (Meng and Takeichi, 2009). In adherens junctions, cadherins form homophilic (more rarely heterophilic), calcium dependent interactions with other cadherin molecules from neighboring cells. The cytoplasmic domain of cadherins is highly conserved among metazoans, distinguishing classical cadherins from other cadherin subfamilies (Hulpiau and van Roy, 2011; Oda and Takeichi, 2011). It contains β-catenin and p120 binding sites, which connects catenins with the actin cytoskeleton in a dynamic manner (Meng and Takeichi, 2009). In comparison with other cadherin subfamilies, classical cadherins are unique by showing the most noticeable variation in their extracellular region among different species (Hulpiau and van Roy, 2011). Indeed, the extracellular domain consists of a variable number of cadherin repeats of about 110 amino-acids each, and - depending on the species - the presence of laminin G and EGF-like domains.

During development, the regulation of expression of specific cadherins correlates with the formation of new tissues. For instance, the folding of the neural tube in vertebrates occurs in parallel with the downregulation of E-cadherin and the upregulation of N-cadherin (Nandadasa et al., 2009). Such cadherin switches are characteristic of several different morphogenetic processes, for example, during gastrulation and neural crest development (Basilicata et al., 2016; Dady et al., 2012; Detrick et al., 1990; Giger and David, 2017; Hatta and Takeichi, 1986; Pla et al., 2001; Rogers et al., 2013; Scarpa et al., 2015; Schäfer et al., 2014; Shoval et al., 2007). During mesoderm formation of *Drosophila*, DE-cadherin becomes replaced by DN-cadherin (Oda et al., 1998), similar to the switch from E- to N-cadherin during mesoderm formation in chicken (Hatta and Takeichi, 1986). It has also been shown that N-cadherin expression triggers active endodermal cell migration, which leads to the cell segregation and germ layer formation (Ninomiya et al., 2012). Moreover, a cadherin switch allows efficient Wnt, BMP and FGF signaling, required for the proper mesoderm differentiation in fruitfly and mouse (Basilicata et al., 2016; Giger and David, 2017; Ninomiya et al., 2012; Schäfer et al., 2014). In line with that, N-cadherin can interact with the FGF receptor and modulate the signaling pathway (Francavilla et al., 2009; Williams et al., 1994). Therefore, accurate control of the expression of cadherin is important for proper cell movements during gastrulation, like epiboly, or convergence and extension of the tissue during axis elongation (Babb and Marrs, 2004; Basilicata et al., 2016; Shimizu et al., 2005; Winklbauer, 2012).

While the role of cadherins has been studied in model bilaterian species, very little is known in diploblastic organisms, such as cnidarians. Most of our knowledge on cell adhesion molecules in cnidarians is restricted to genome analyses (Hulpiau and van Roy, 2009; Hulpiau and van Roy, 2011; Tucker and Adams, 2014). The sea anemone *Nematostella vectensis* has become one of the prime model organisms for embryonic development during the last two decades (Genikhovich and Technau, 2009a; Layden et al., 2016; Technau and Steele, 2011). A bioinformatic analysis of the available genome sequence of *Nematostella vectensis* (Putnam et al., 2007) revealed that it has 16 different cadherins from all main groups of the cadherin superfamily (classical, flamingo, FAT, dachsous, FAT-like, protocadherins and cadherin-related proteins) (Hulpiau and van Roy, 2011). It has been shown that adherens junctions in *Nematostella* ultrastucturally resemble those in bilaterians (Fritzenwanker et al., 2007). However, up to date, the molecular composition of these junctions has not been described, although a recent report questioned the presence of adherens junctions in the inner layer of *Nematostella* (Salinas-Saavedra et al., 2018).

Germ layers are formed in *Nematostella* by invagination of the endoderm at the animal pole (Kraus and Technau, 2006; Magie et al., 2007). However, whether classical cadherins play a role in germ layer formation of a non-bilaterian is not known. Here, we show that classical cadherins of *Nematostella*, Cadherin1 (Cdh1) and Cadherin3 (Cdh3), form the adherens junctions of the epithelium of both germ layers. Germ layer differentiation in *Nematostella* is marked by a cadherin switch, where Cdh3 is downregulated in the inner, endodermal layer, while Cdh1 is upregulated and remains the only cadherin expressed in the endoderm. Unexpectedly, we found that in addition to the apical adherens junctions both Cdh1 and Cdh3 are also involved in cell junctions between cells on the basal side. Knockdown experiments of *cdh1* and *cdh3* indicate important roles of cadherins in cell adhesion and tissue morphogenesis of this diploblastic metazoan.

## Result

### Structure of classical cadherins of *Nematostella vectensis*

Three genes encoding classical cadherins have been predicted in the genome of *Nematostella vectensis – cadherin1, cadherin2, cadherin3 (cdh1, cdh2, cdh3)*. All three genes are the result of lineage-specific duplications (Hulpiau and van Roy, 2011). Expression of *cdh2* is not supported by the RNAseq data, and it is not detectable by *in situ* hybridization. Therefore it might be a pseudogene and it was not investigated here.

These predictions suggested a structure of the classical cadherins, which would be composed of a typical intracellular domain with binding sites for ß-catenin and p-120, and a large extracellular domain consisting of three EGF-like and two Laminin G domains, followed by 32 or 25 extracellular (EC) repeats (Fig. 1A), while a fruit fly has 17 EC repeats, chick - 13 EC repeats, mouse only 5 EC repeats (Hulpiau and van Roy, 2011). This would result in an mRNA of >; 15 kb and predict a protein size of about 480 kDalton. However, in a recent publication, two cadherins (called Cadherin 1 and Cadherin 2) with 14 and 30 Cadherin repeats, respectively, were reported (Clarke et al., 2016). To resolve this discrepancy, we used numerous primers derived from the predictions of Hulpiau and van Roy and our gene models from RNAseq (Fredman et al., 2013), cloned the large transcript in overlapping pieces of about 2-3 kb. Our results show that both cdh1 and cdh3 possess 31 CA repeats, which is closer to the original model of Hulpiau and van Roy (Fig. 1A). We therefore follow their gene terminology. Thus, the classical cadherin genes encode huge proteins, compared to the bilaterian counterpart, leading to the question, of whether they play similar or different roles in the development of *Nematostella*.

**Fig. 1.**
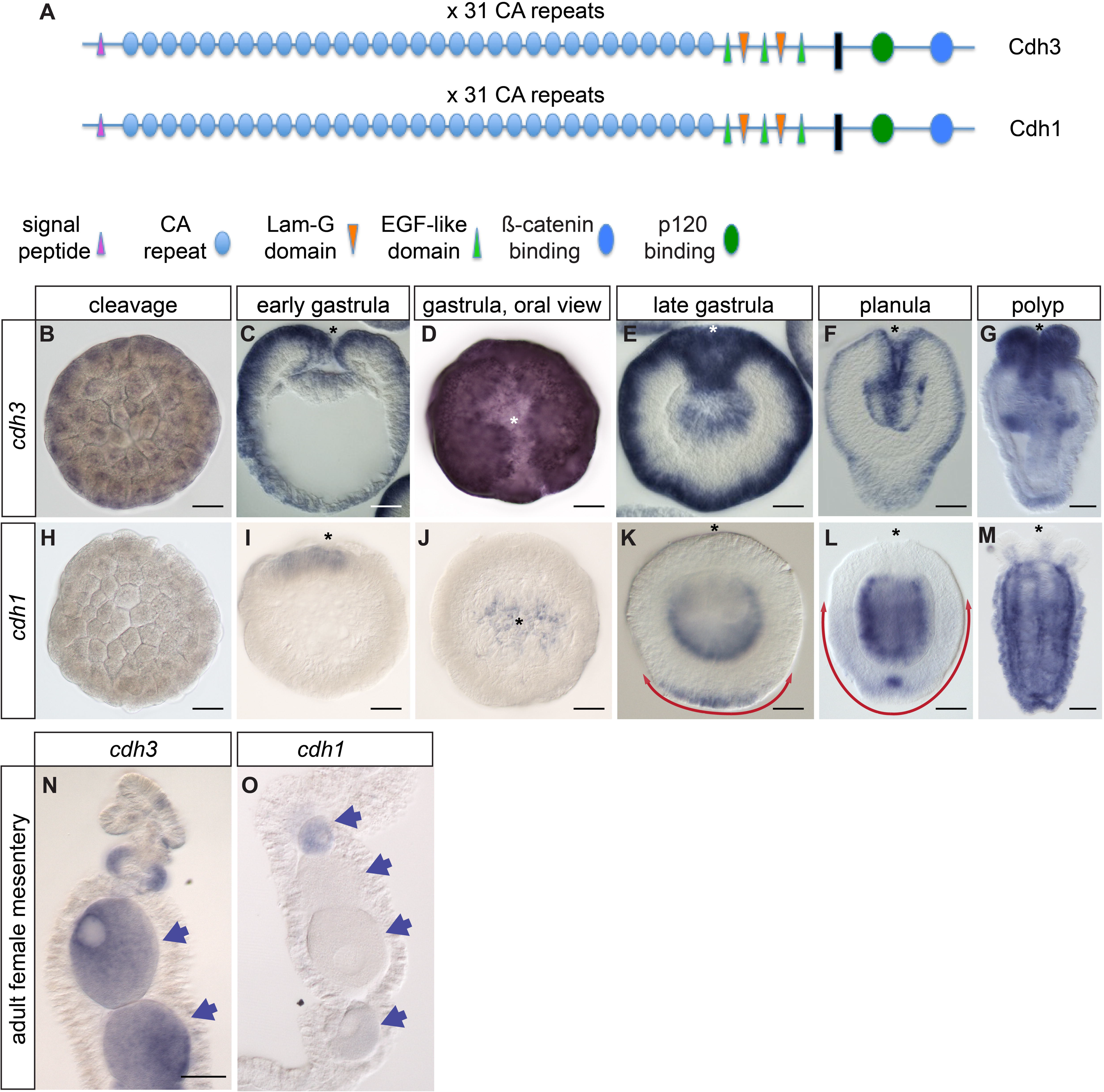
*cadherin3* and *cadherin1* expression is highly dynamic during early development and polyp growth. (A) Protein domain organization of classical cadherins of *Nematostella vectensis*. (B, H) Cleavage. (C, I) Early gastrula, lateral section. D, J: Early gastrula, oral view. E, K: Late gastrula, lateral section. (F, L) Planula, lateral section. (G, M) Primary polyp. (N, O) Adult mesentery section. Double-headed arrows show an expansion of *cdh1* expression on the aboral pole. Arrows show the eggs. Asterisk indicates an oral pole. Scale bar (A-M) 50 μm; (N-O) 100 μm.

### Expression of classical cadherins is highly dynamic during early development of *Nematostella*

To characterize the pattern of classical cadherins gene expression during normal development, *in situ* hybridization has been performed on all developmental stages, starting from early cleavage till adult polyp. *cdh3* is maternally expressed at significant levels, which are detectable at the earliest cleavage stages. *cdh3* is then strongly expressed in all cells starting from the egg until gastrula stage (Fig. 1B-E,N). During early gastrulation *cdh3* expression is decreased in the presumptive endoderm (Fig. 1C,D). Later on, *cdh3* is completely down-regulated in the endoderm at planula stage (Fig. 1C-F).

By comparison, *cdh1* expression could not be detected by in situ hybridization until early gastrula stage (Fig. 1H-J), although RNAseq data suggest that it also maternally expressed at lower levels (Casper et al., 2018). During gastrulation *cdh1* expression first appears and intensifies in the pre-endodermal plate (Fig. 1I,J). At late gastrula stage *cdh1* starts being expressed in the aboral ectoderm and expands then orally during planula development (Fig. 1 K,L). Interestingly, at the late planula stage, the strongest *cdh1* expression is detected in the endoderm and in the subpopulation of ectodermal cells which give rise to a sensory apical organ (Fig. 1L).

In primary polyp, *cdh3* expression remains strongly expressed in the tentacles and in the pharynx and weakly in the body-wall ectoderm (Fig. 1G). Almost complementary to that, *cdh1* is expressed both in the ectoderm and endoderm, but it is completely excluded from the ectoderm of the pharynx and tentacles (Fig. 1M). In juveniles, *cdh1* is expressed in the endoderm and body-wall ectoderm, but not in the ectoderm of the pharynx. Interestingly, the part of the pharynx carrying siphonoglyph and the ciliary tract below the pharynx specifically expresses *cdh1* (Fig. S1C-H). In adults, *cdh1* is expressed in the body-wall endoderm and in small oogonia (Fig. 1O; Fig. S1I). Interestingly, *cdh3* in juveniles and adults is detectable only in the ectoderm of the pharynx and tentacles, ciliated tract, septal filaments, with no expression in the body-wall ectoderm (Fig. S1A,B).

### Cdh3 is the main component of adherens junctions during cleavage and gastrulation

We wished to visualize the subcellular localization of Cdh3 protein during development. We therefore expressed fragments of the Cdh1 and Cdh3 proteins and generated specific polyclonal and monoclonal antibodies against Cdh1 and Cdh3 respectively (Materials and Methods). Immunocytochemistry experiments were carried out in all stages of development. Cdh3 protein is already accumulated at the apical cell junctions at the first cell divisions, suggesting an unappreciated early cell polarity. It is also detectable in less confined areas at the lateral contacts between cells (Fig. 2A-C). Interestingly, cells maintain their polarity during cell divisions. Cdh3 stays localized at the apical cell junctions at different cell cycle stages (Fig. 3). In contrast to the Par system proteins (Ragkousi et al., 2017; Salinas-Saavedra et al., 2018) we did not observe a loss of cell polarity marked by Cdh3 during early cell divisions (Fig. 3). Later, during blastoderm formation, apical cell junctions become more pronounced (Fig. 2D-F). Strikingly, we found that Cdh3 is localized not only on the apical-lateral side but also on the basal-lateral side of the cells (Fig. 2D-L). Notably, the ultrastructural analysis by transmission electron microscopy revealed that the cell-cell junctions at the basal side resemble the adherens junctions at the apical side (Fig. 2M,N). Thus, *Nematostella* has a unique epithelium, where cells form cell-cell contacts on both apical and basal sides. These Cdh3-positive junctions develop before any contact to an endodermal layer or presence of the mesoglea, the extracellular matrix of Cnidaria. This is remarkable, and has to our knowledge not been described in any other animal. Interestingly, as the pre-endodermal plate (PEP) starts to invaginate and the cells adopt a partial EMT phenotype, Cdh3 disappears from the basal junctions of the invaginating cells (Fig. 2G,H,I). Meantime, ectodermal cells of the blastoderm retain both apical and basal cell contacts. As the pre-endodermal cells lose basal junctions, its epithelium becomes less rigid and columnar. Pre-endodermal cells form filopodia and become more motile on the basal side (Fig. 2O). This event is possibly one of the crucial steps of the incomplete EMT, which pre-endodermal cells undergo during gastrulation (Kraus and Technau, 2006; Salinas-Saavedra et al., 2018). Notably, apical cell junctions expressing Cdh3 are preserved in the pre-endodermal cells during the course of gastrulation (Fig. 2J-L).

**Fig. 2.**
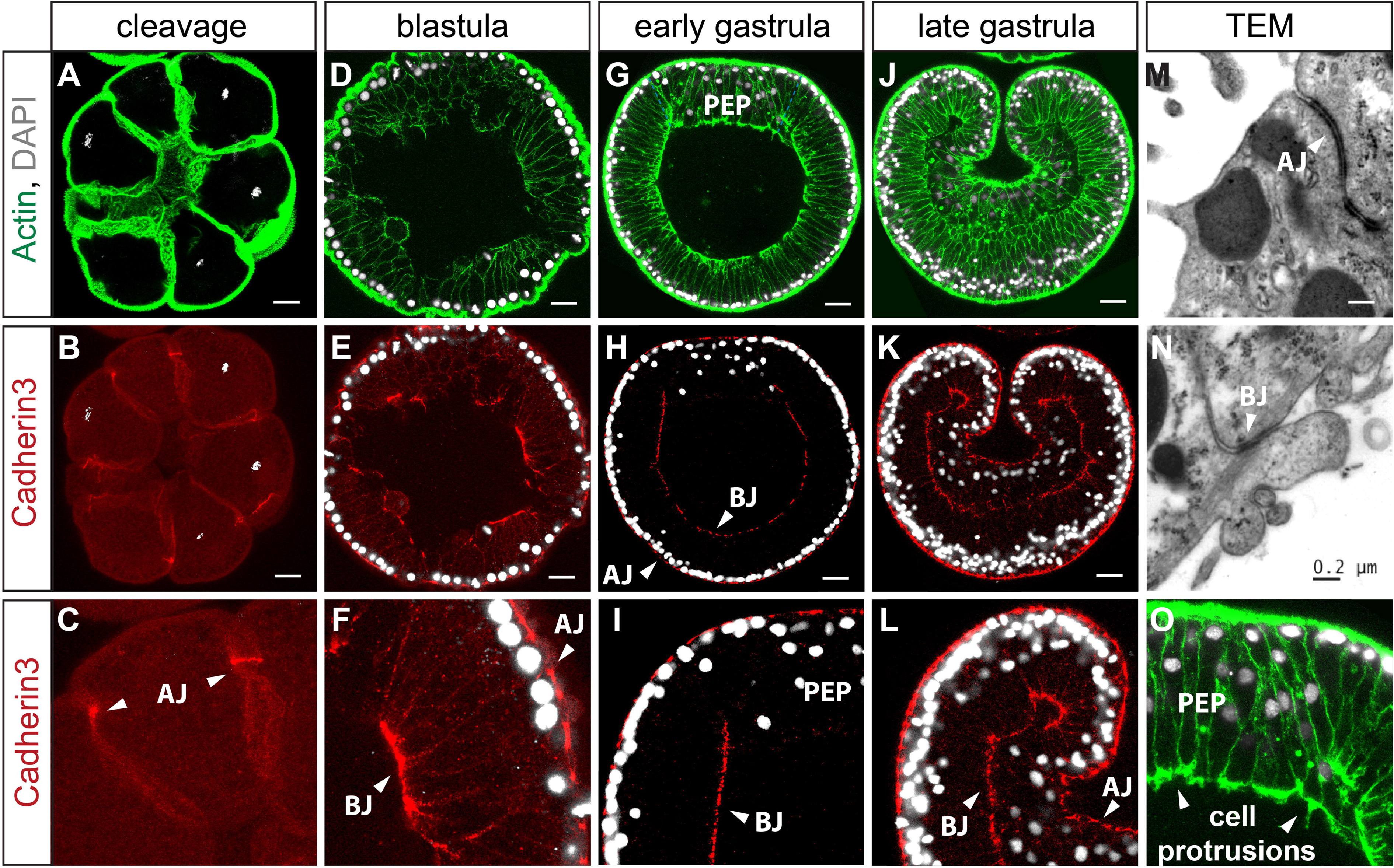
Cdh3 is a major component of adhesion complexes during cleavage and gastrulation. (A-F) Besides apical adherens junctions (AJ), strong basal epithelial contacts (BJ) form in the blastula during epithelialization. (G-I) As the pre-endodermal plate (PEP) starts to invaginate, Cdh3 disappears from the basal junctions and decreases in the apical junctions in the pre-endodermal plate. Ectodermal cells preserve both apical and basal cell contacts. (J-L) Late gastrula. Apical junctions are present in the ectoderm and in the endoderm. (M-N) Transmission electron microscopy (TEM) of the *Nematostella* epithelium. (O) Cell protrusions on the basal side of the PEP. Apical cell-cell junctions (AJ); basal cell-cell junctions (BJ). Scale bar 20 μm; Scale bar M-N: 0,2 μm.

**Fig. 3.**
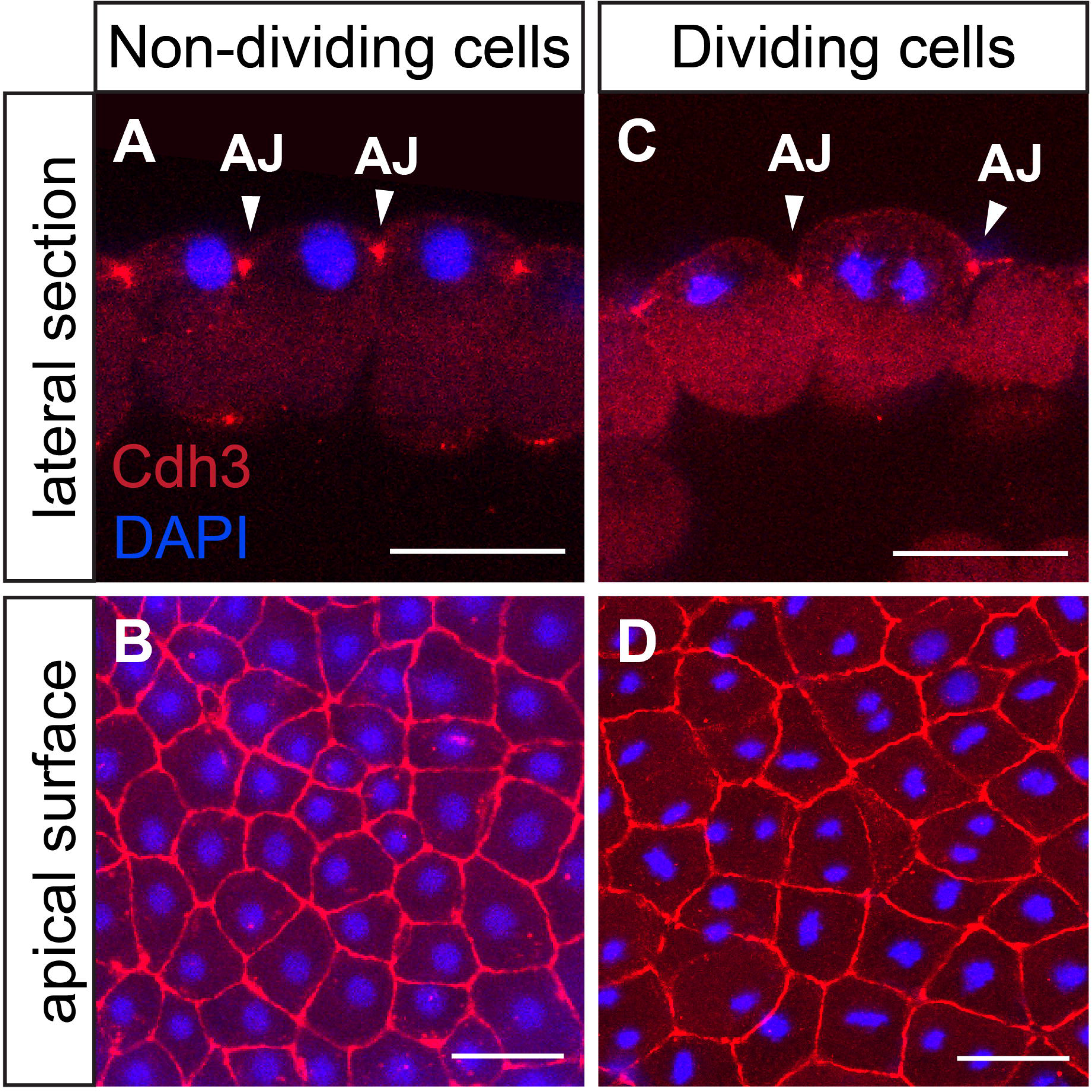
Cdh3 apical junctions localization and cell polarity are preserved during the cell division. (A, B) Non-dividing blastula cells. (C, D) Dividing blastula cells at different mitotic phases. Scale bar 25 μm.

After invagination is completed, Cdh3 fully disappears from the cell junctions of the endoderm, in line with the decrease of mRNA expression in the whole endoderm. Cdh3 remains expressed exclusively in the ectoderm, forming apical and basal adherens junctions (Fig. 4A-E). Notably, while the boundary between ectoderm and endoderm is very difficult to discern by morphological criteria, Cdh3 localization at the cell junctions in the pharynx precisely marks the boundary between the last ectodermal and the first endodermal cell (Fig. 4B,E). At polyp stage Cdh3 remains exclusively expressed in the ectoderm, with especially strong expression in the pharynx and the tentacles (Fig. 4F,G).

**Fig. 4.**
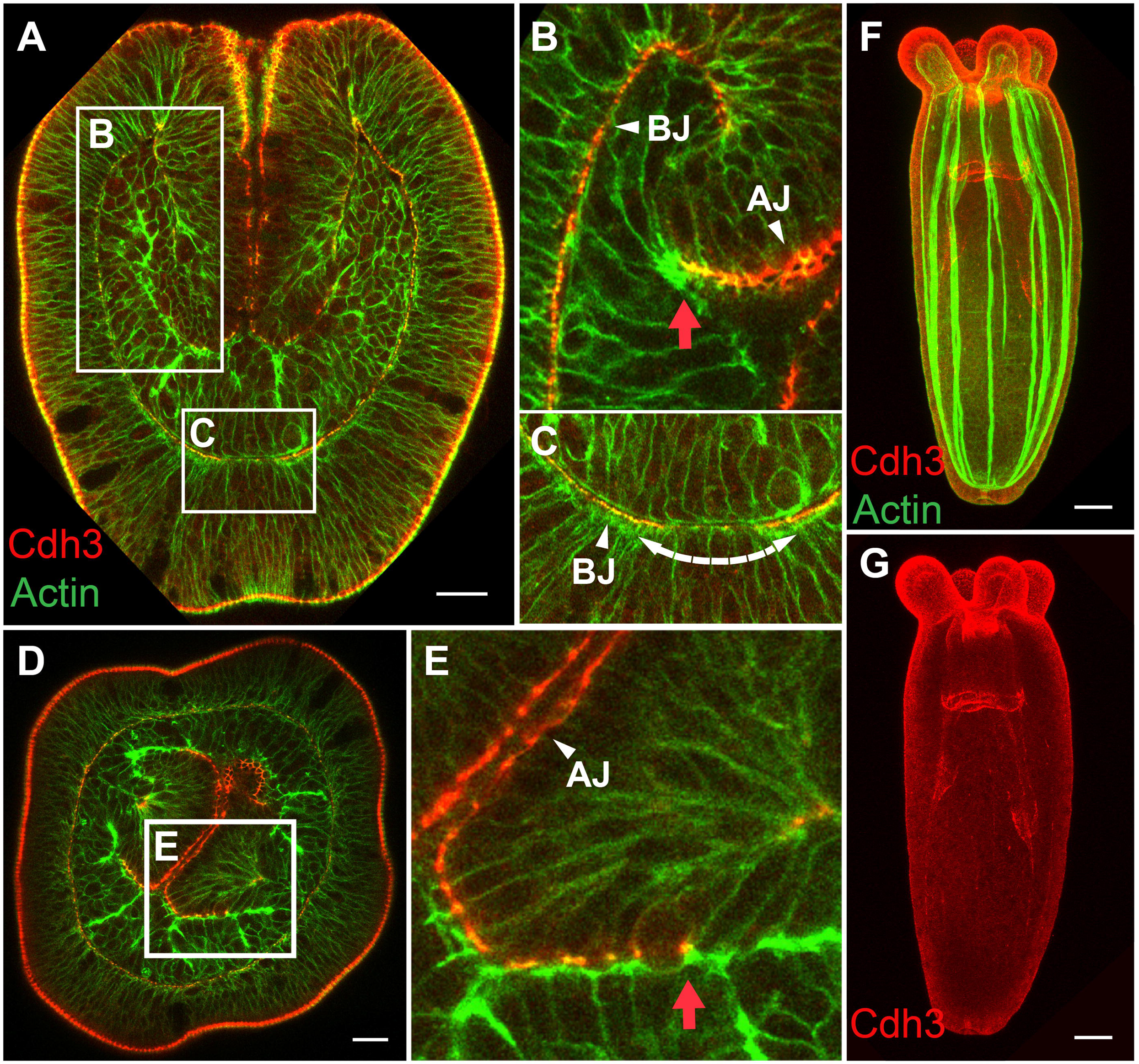
Cdh3 marks the boundary between ectoderm and endoderm. (A-C) Lateral section of planula. (D, E) Cross-section of planula. Ectodermal-endodermal boundary in the pharynx is distinctly labeled by Cdh3 localization in the cell junctions. (F, G) Primary polyp. Cdh3 is expressed exclusively in the ectoderm, forming apical and basal adherens junctions. Red arrow indicates the boundary between the last ectodermal and the first endodermal cell. Scale bar 20 μm.

### Cdh1 protein expression marks a cadherin switch during endoderm formation

After completion of gastrulation Cdh1 protein forms pronounced cellular junctions. At early planula larva Cdh1 localizes to the apical and basal junctions of the endoderm (Fig. 5A-D). Hence, the formation of the endoderm is marked by a cadherin switch from Cdh3 to Cdh1.

**Fig. 5.**
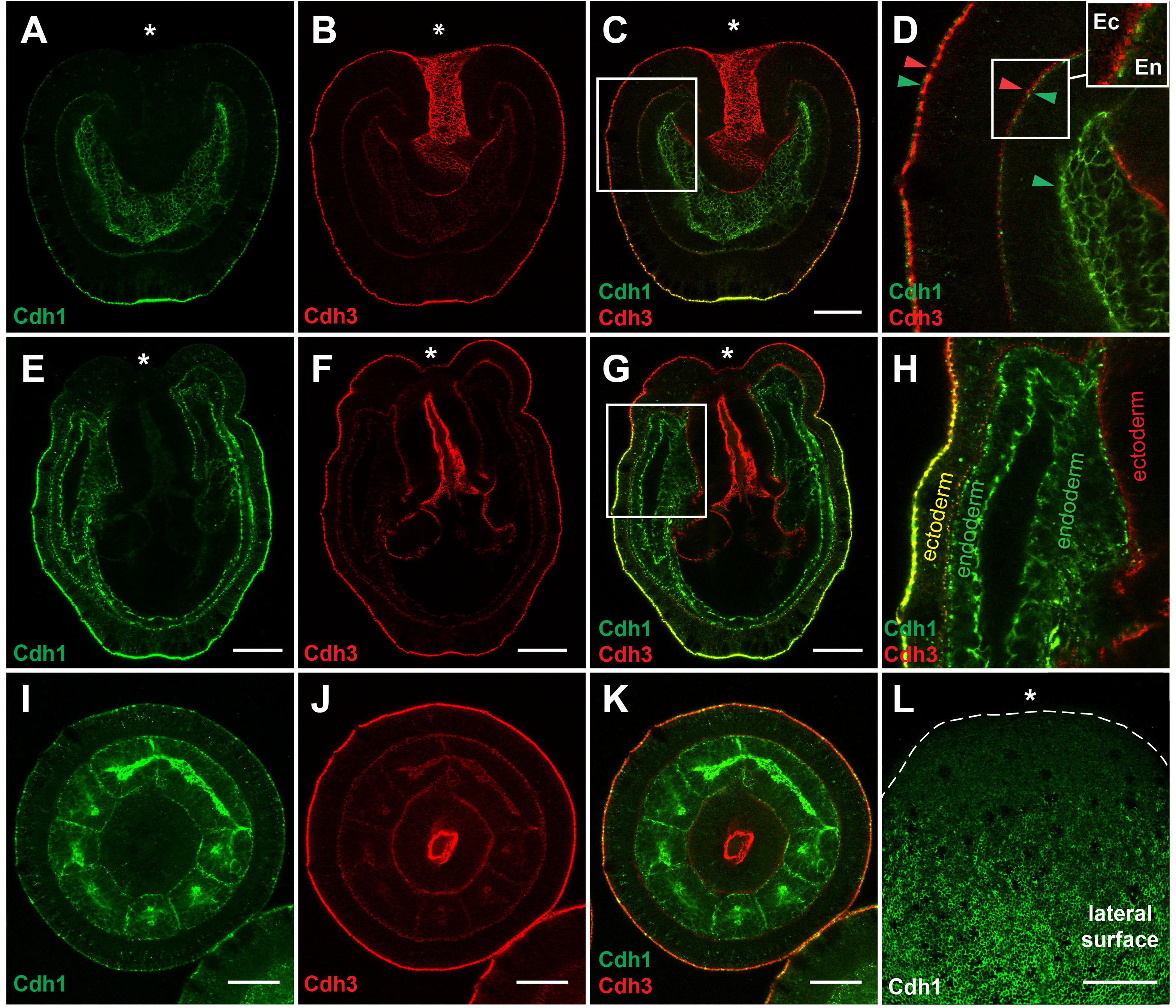
Cdh1 and Cdh3 protein localization during germ layer differentiation. (A-D) Planula lateral section. (E-H) Lateral lection of the primary polyp. (I-K) Planula cross-section. (L) Surface of the planula, oral part of the ectoderm is free of Cdh1. Cdh1 is localized in the apical and basal junctions of the endoderm, as well as in the apical junctions and basal junctions of the aboral ectoderm, especially in the area of the apical organ. Cdh1 is gradually disappearing from the ectoderm towards the oral pole and completely excluded from the ectoderm of the tentacles and the pharynx Cdh3 is localized to the apical and basal junctions of the body wall ectoderm, ectoderm of the pharynx and is completely excluded from the endoderm. Asterisk marks an oral pole. Scale bar 50 μm.

In addition to the endodermal expression, Cdh1 starts being strongly expressed in the apical organ ectoderm and then expands into a wider domain in the aboral ectoderm, where Cdh1 and Cdh3 are co-expressed (Fig. 5). At the ectodermal surface, expression of the Cdh1 decreases in a gradient manner towards the oral pole (Fig. 5L). Interestingly, the ectodermal cell population, which gives rise to the apical tuft, is also different from the rest of the ectoderm in terms of cadherin expression. These cells lose Cdh3 basal junctions, while keeping the apical junctions (Fig. 4C). We assume that the loss of the basal junctions is due to a different anchoring of the cells to the mesogloea. This specific arrangement may be connected with the special function of these cells (Fig. S2). Interestingly, the loss of Cdh3 expression in the ectodermal apical tuft cells goes hand in hand with an upregulation of Cdh1 in these cells (Fig. 1L; Fig. 5A,C).

Thus, in summary, Cdh3 is the major component of adhesion complexes during cleavage and gastrulation and it is present in all cells till late gastrula stage. Cdh3 forms apical and basal cell adherens junctions in the blastodermal epithelium. During invagination of the pre-endodermal plate basal cell junctions disappear from the future endoderm. Further endoderm differentiation leads to the complete Cdh3 to Cdh1 replacement. Therefore, there is a distinct boundary between ectoderm and endoderm which is defined by Cdh1 / Cdh3 localization.

### Cdh3 in apical ectodermal adherens junctions colocalizes with β-catenin

A recent biochemical study showed that the intracellular domain of the classical cadherins can form a ternary complex with α-catenin and β-catenin (Clarke et al., 2016). To explore further the molecular composition of the adherens junctions, we co-stained the embryos with the antibody against Cdh3 and β-catenin. At blastula stage Cdh3 is co-localized with ß-catenin at the apical junctions whereas basal junctions do not show such pronounced co-localization (Fig. 6A-C; G-I). Interestingly, at planula stage β-catenin was detected only in the apical junctions of the body wall ectoderm but neither in the ectodermal pharynx nor the endoderm (Fig. 6D-F). These results could mean that not all the cell contacts of *Nematostella* epithelium contain β-catenin, in line with other recent findings (Salinas-Saavedra et al., 2018). This is surprising, as no ultrastructural differences in the junctions of endoderm and ectoderm could be detected (Fig. S4).

**Fig. 6.**
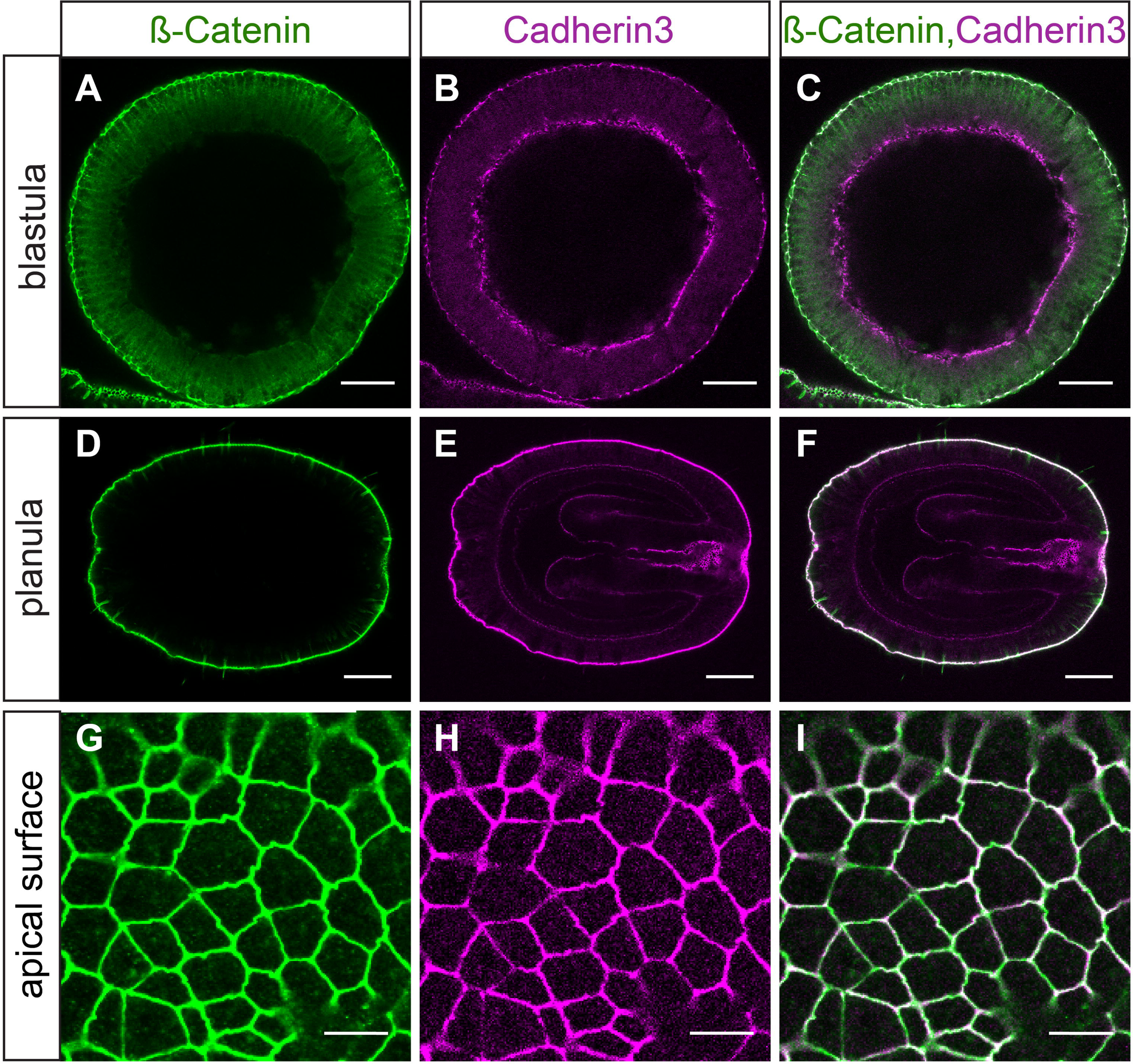
Cdh3 and beta-catenin are co-localized at the apical (and partially at the basal) cell junctions of the ectoderm. (A-C) blastula stage. (D-F) planula stage. (G-I) Apical surface of the ectoderm. Scale bar (A-F) 50 μm; (G-I) 10 μm.

### Function of classical cadherins in early development

To examine the function of cadherins we performed knockdown experiments using morpholinos and shRNA. First, we injected independently two non-overlapping translation blocking *cdh3* morpholinos (MO). However, we could still detect Cdh3 in apical and basal cell junctions in the whole mount MO injected embryos (Fig. 7). Indeed, on the ultrastructural level, the adherens cell junctions looked similar in morphants and in control embryos (Fig. 7C,F). These results can be explained by the significant maternal deposition mRNA and protein. However, development of Cdh3 morphants was arrested after gastrula stage, presumably due to the block of translation of zygotically expressed *cdh3*. As a result, when Cdh3 protein becomes limited, post-gastrula embryos are unable to develop further (Fig. 7A,B).

**Fig. 7.**
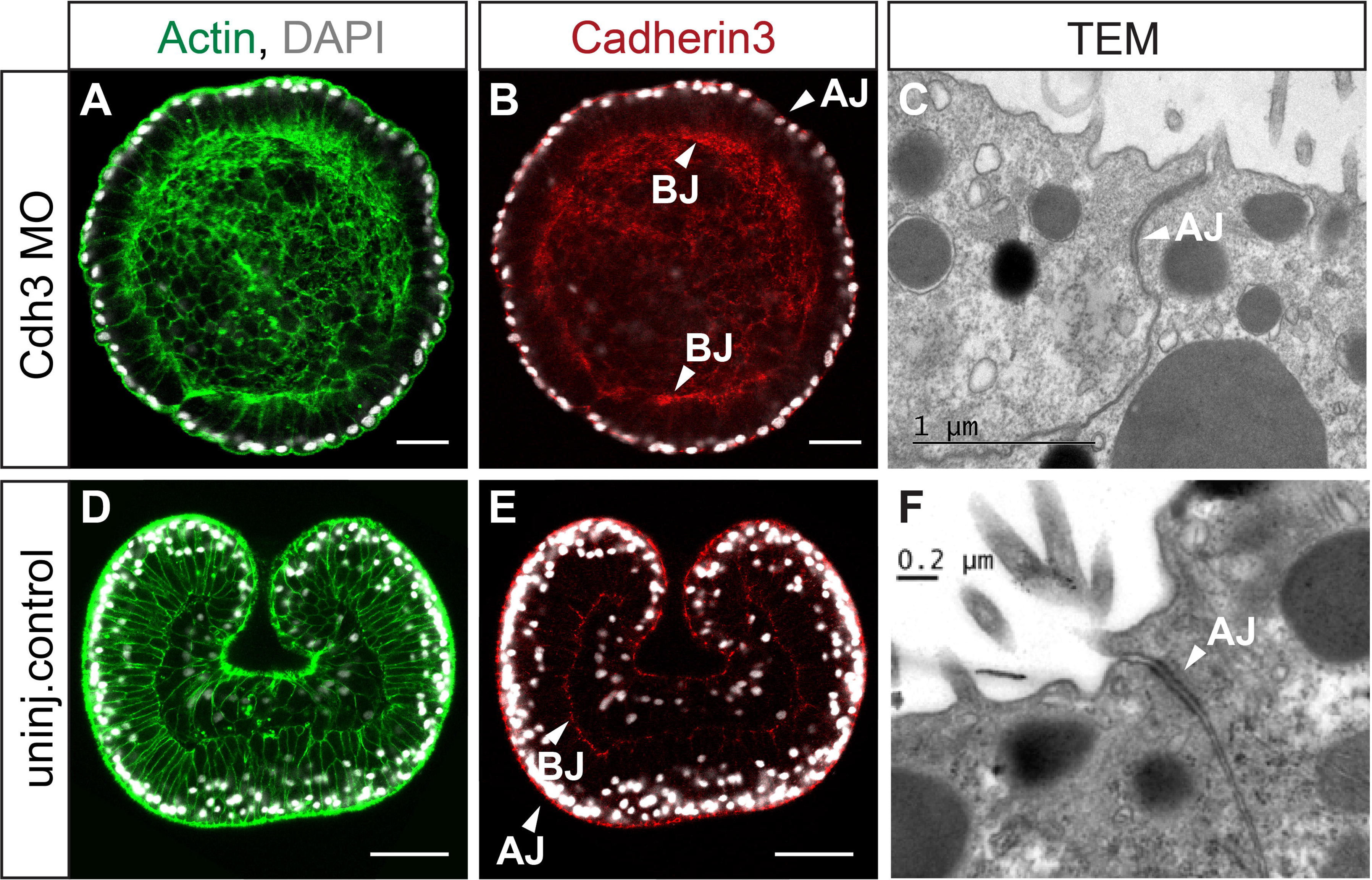
Cdh3 knockdown blocks gastrulation movements. (A-C) Cdh3 morpholino (MO) injected embryos, 28 hours post-fertilization (hpf). (D-F) Control embryos, 28 hpf. Apical (AJ) and basal (BJ) cell junctions of Cdh3 morphants look very similar to the cell junctions of the control gastrulae. Scale bar (A,B,D,E) 40 μm.

The mild knockdown effect on the presence of Cdh3 in the junctions also suggests that there is relatively little turnover in established junctions. Therefore, to assess the function of Cdh3 in establishing new cell junctions we used an aggregate assay. *Nematostella* gastrulae can be dissociated into single cells and small clusters and can be re-aggregated by centrifugation into the cell aggregates (Kirillova et al., 2018). We followed the establishment of cell contacts and the formation of the epithelium in the developing cell aggregates (Fig. 8). Dissociated cells do not show any signs of polarization: Cdh3 is not localized to any side (Fig. 8C). Only 30 min after re-aggregation, Cdh3 becomes localized to the apical side of the outer cells of the aggregate (Fig. 8E,F,M). The first signs of epithelialization become apparent. 12 hours after re-aggregation the outer epithelial layer is completely formed and Cdh3 is localized at the apical and basal cell junctions (Fig. 8H,I). 24 hours after re-aggregation two epithelial layers - ectoderm and endoderm are formed. Both of the layers possess basal and apical cadherin cell junctions (Fig. 8K,L,S). Cdh1 starts being expressed in both ectoderm and endoderm at 24 hours of aggregate development (Fig. 8N,Q,T,W). Similar to the normal embryo, ectoderm expresses both Cdh1 and Cdh3, while endoderm expresses exclusively Cdh1 (Fig. 8U,X).

**Fig. 8.**
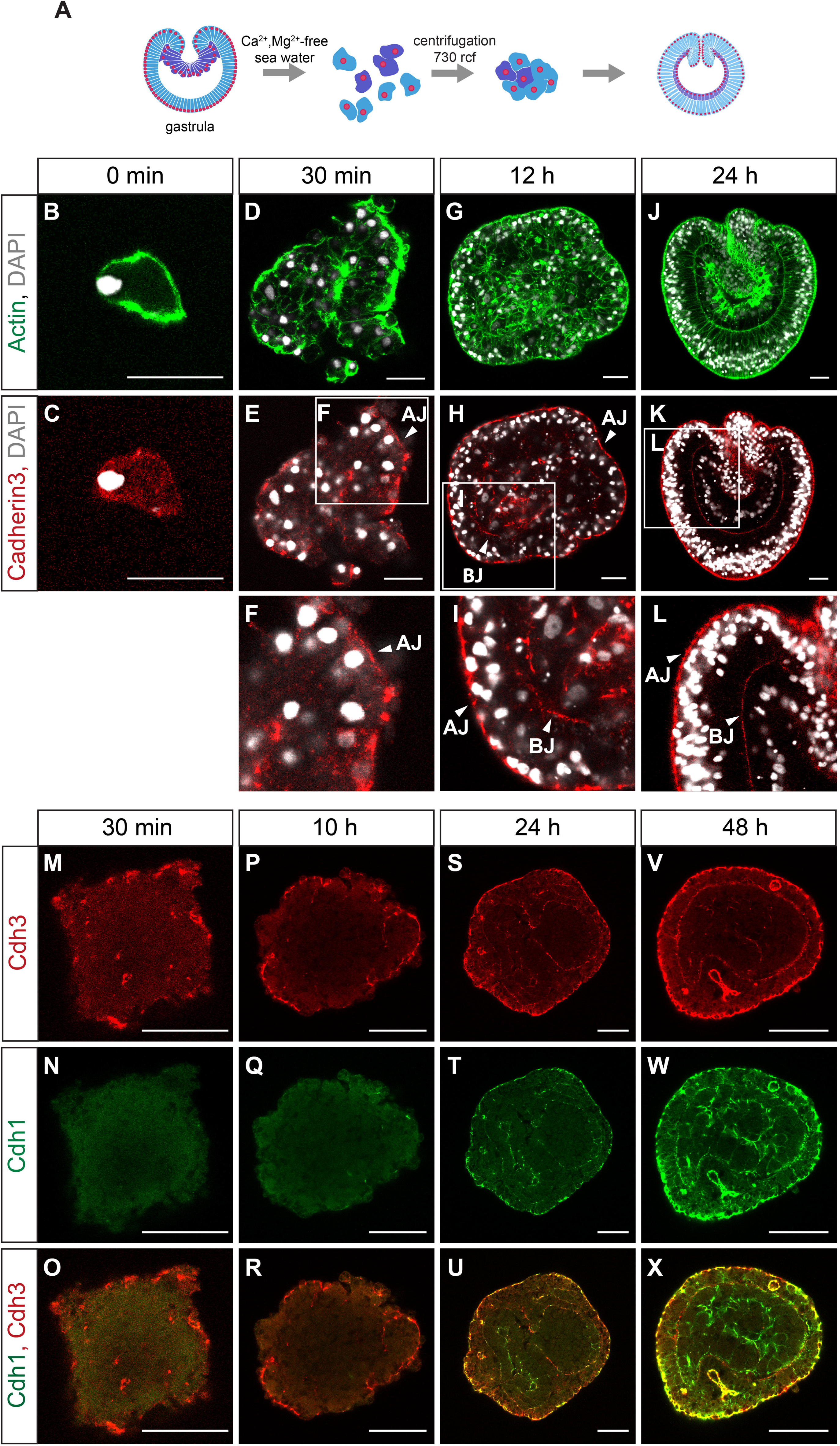
Reestablishment of the polarity and de novo germ layer formation in the cell aggregate. (A) Scheme of the experiment. (B,C) Dissociated cells do not show polarized Cdh3 localization. (D-F) Epithelialization of the cell aggregate starts ~30 min after re-aggregation. (G-I) 12h after dissociation ectoderm of the aggregate is fully epithelialized. (J-L) 24h. Aggregate forms two germ layers. (M-X) Cdh1 protein appears at the junctions at 24h of aggregate development. At 48h after re-aggregation Cdh1 is broadly expressed in both germ layers. Apical junctions (AJ); Basal junctions (BJ). Scale bar (B-K) 20 μm; (M-X) 50 μm.

To address the question how Cdh3 down-regulation influences the establishment of new cell contacts in the aggregate, we dissociated an equal amount of *cdh3* MO injected gastrulae and standard MO injected gastrulae as a control. The first difference we observed was that the size of the aggregates from *cdh3* morphant cells was significantly smaller than control aggregates (p < 0.0001) (Fig. 9K-M). Moreover, aggregates from *cdh3* morpholino injected embryos start falling apart into cells immediately after re-aggregation (Fig. 9; Movies 1,2). Ultrastructural imaging with TEM confirmed that cells in Cdh3 MO aggregates do not form well-defined subapical adherens junctions, while in the control aggregate cell contacts are well developed (Fig. 9E,J). Interestingly, cells in the Cdh3 MO aggregates make lamella-like protrusions extending to the neighboring cell on the apical surface (Fig. 9E). These results show that *cdh3* knockdown impairs the *de novo* formation of the cell contacts, although it does not affect the earlier established contacts built from the maternal protein.

**Fig. 9.**
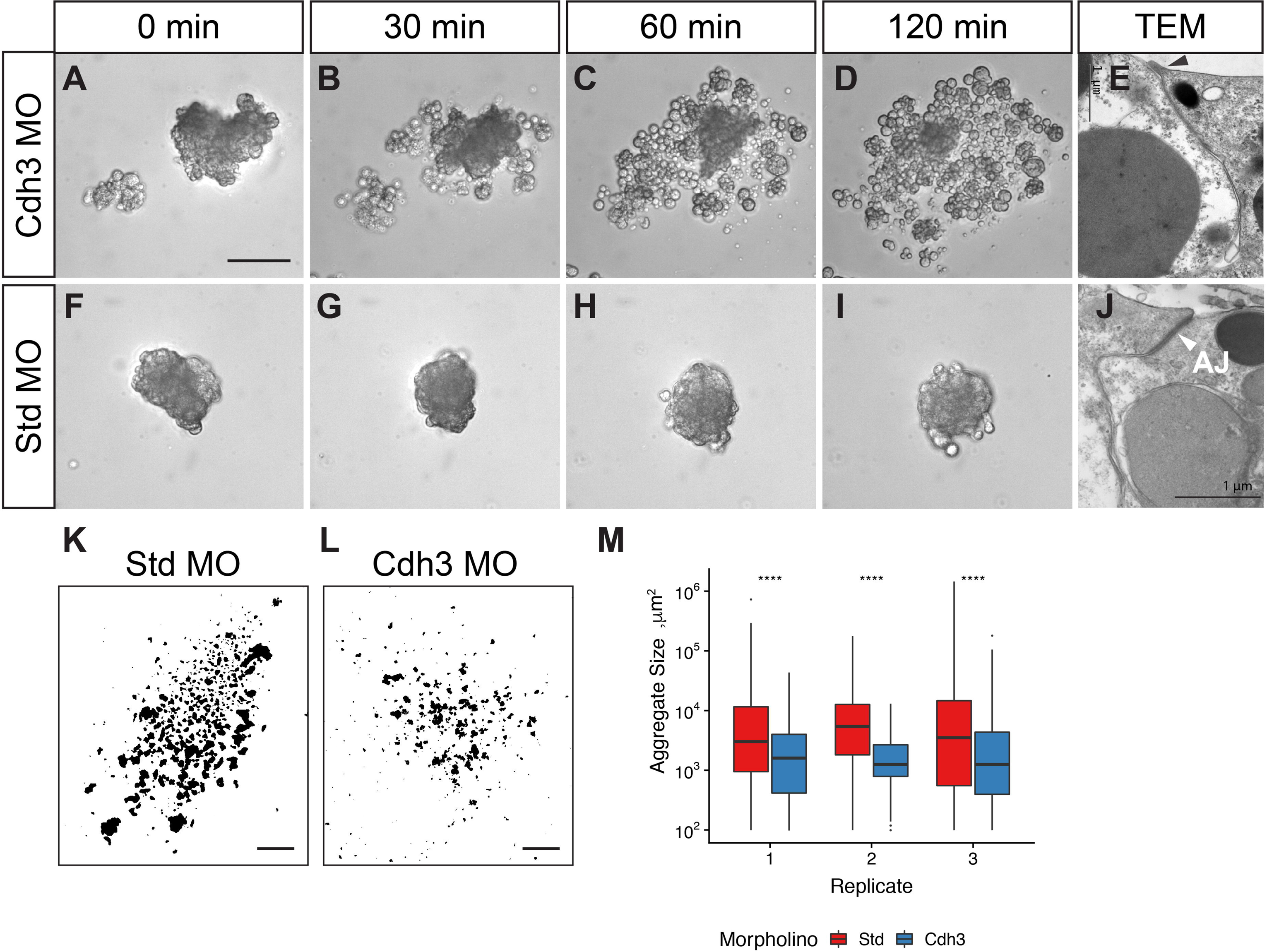
Cdh3 MO aggregates fail to form adherens junctions *de novo*. (A-D) Cdh3 morpholino (MO) aggregates do not form new cell contacts, fail to develop and fall apart into cells. (F-I) Standard (Std) MO control aggregates stay compact. (E,J) TEM of the apical adherens junctions. Apical cell junctions (AJ) of the Cdh3 MO aggregates are much less pronounced than AJ in the control aggregates. (K-M) Cdh3 MO aggregates are significantly smaller than Std MO aggregates. Distribution means within each replicate were tested for significance using a two-sided unpaired Wilcoxon rank-sum test (****: p < 0.0001). Scale bar (A-D; F-I) 50 μm; (E, J) 1 μm. (K-L) 1 mm.

To further explore the role of Cdh1 protein, we down-regulated *cdh1* using an independent approaches, the shRNA-mediated knockdown (He et al., 2018). As in MO knockdown, shRNA knockdown leads to the significant decrease of the Cdh1 protein as assayed by immunohistochemistry (Fig. S3; Fig. 10). Although early development including gastrulation appears largely unaffected, in the subsequent planula stage, mesenteries do not form upon *cdh1* knockdown. In all MO and shRNA injected embryos mesenteries were absent or impaired, while there are eight mesenteries developed in the control embryo at this stage (Fig. 10; Fig. 11C,F). Besides the predominant expression in the endoderm, *cdh1* is also expressed in the apical tuft region of the ectoderm (Fig. 1L). Interestingly, *cdh1* knockdown abolishes expression of FGFa1, which is responsible for the apical organ development (Rentzsch et al., 2008). In most of the *chd1* MO injected embryos the apical organ does not form and there is lack of FGFa1 expression (Fig. 11). These results suggest that Cdh1 is crucial for morphogenesis and differentiation of the endoderm as well as the development of the apical organ.

**Fig. 10.**
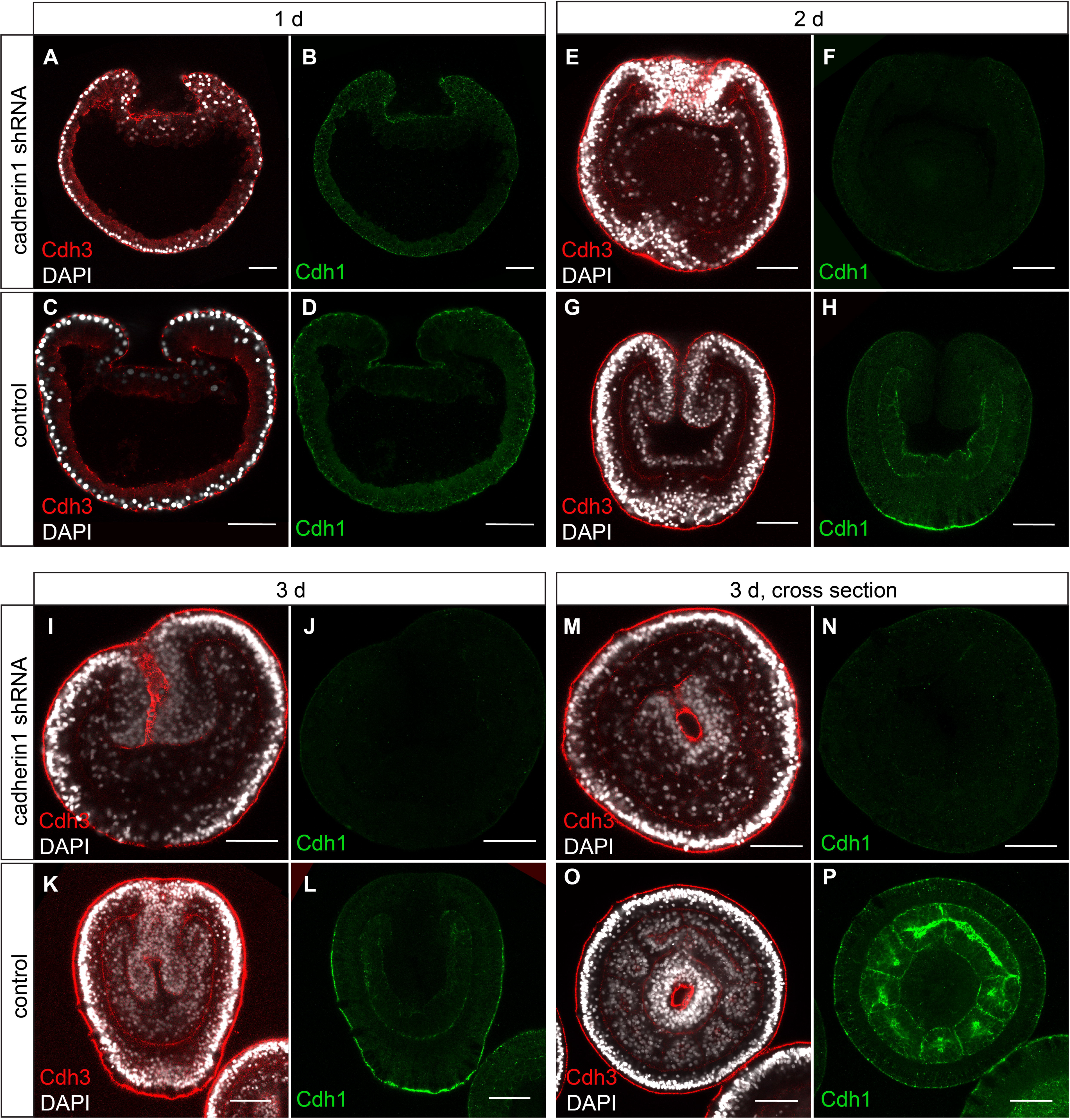
Mesenteries do not develop upon Cdh1 knock-down by shRNA injection. Cdh1 protein expression is strongly down regulated. (A-D) 1 dpf, gastrula stage, lateral section. (E-H) 2dpf planula, lateral section. (I-L) 3 dpf planula, lateral section. (M-P) 3d planula, cross section.

**Fig. 11.**
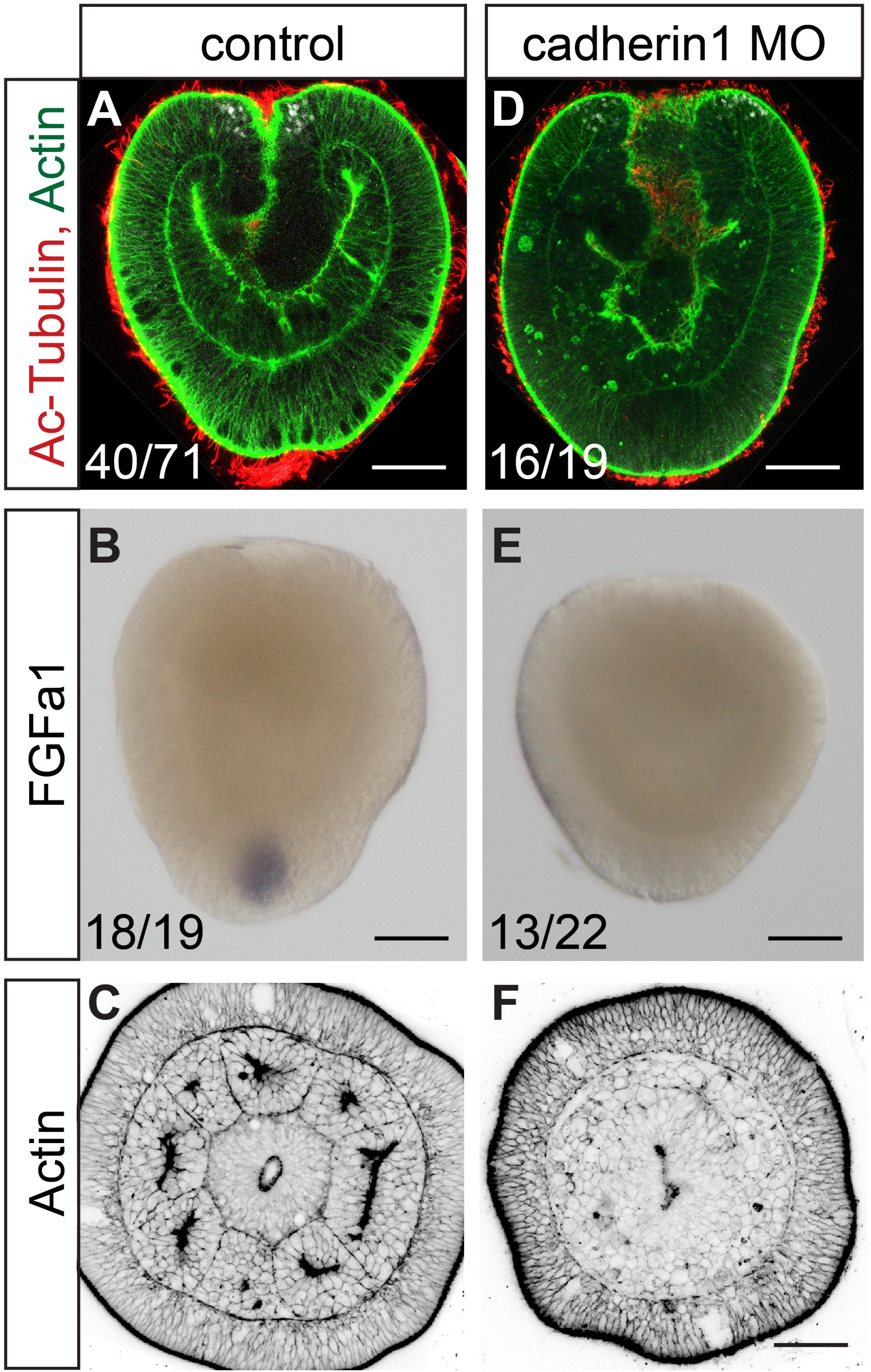
Cdh1 knock-down impairs apical organ development. (A-C) Control embryo. (D-F) Cdh1 MO knockdown. Apical organ fail to develop (acetylated tubulin antibody staining). FGFa1 is not expressed. Mesenteries do not form (phalloidin staining). Scale bar 50 μm.

## Discussion

### Evolution and structure of cadherins

Although proteins with cadherin domains are present in choanoflagellates, cadherins with intracellular catenin binding domains are an important class of cell adhesion molecules that arose only in metazoans (Nichols et al., 2012). They are mediating not only cell adhesion between epithelial cells, but are strongly involved in the differentiation of specific cell types. Recently, they have been shown to also convey mechanotransduction, i.e. the activation of gene expression via translocation of β-catenin to the nucleus upon mechanical stress (Pukhlyakova et al., 2018; Röper et al., 2018). However, most studies on the role of cadherins have been carried out in bilaterian model organisms like mouse or *Drosophila*. Here, we show for the first time the localization and function of both classical cadherins in a representative of the Cnidaria, the sea anemone *Nematostella vectensis*. The two investigated *cadherin* genes code for large proteins with 31 EC domains each, largely confirming previous predictions from the genome (Hulpiau and van Roy, 2011) and gene models based on our transcriptome assembly (Fredman et al., 2013). This significantly extends the structure of the recently published gene model for Cdh3 (termed Cad1 in (Clarke et al., 2016). Thus, cnidarians as well as other non-bilaterians have substantially larger classical cadherins than most bilaterians and their extracellular domain structure is reminiscent of the FAT-like proteins (Hulpiau and van Roy, 2009; Hulpiau and van Roy, 2011). It will be interesting to determine which extracellular domains are engaged in homophilic or heterophilic interactions.

### Cadherins are localized to apical and basal junction in both germ layers

Both cadherins are localized to cell-cell junctions in the epithelial cells of both ectoderm and endoderm (Fig. 12A). Of note, both cadherins do not only localize to apical junctions but also to basal cell-cell junctions. Electron and confocal microscopy analyses suggest that these are adherens junctions. This is in contrast to a recent study, which claimed that the endodermal epithelium does not contain adherens junctions, since neither Par complex components nor β-catenin could be detected (Salinas-Saavedra et al., 2018). Yet, we could also detect β-catenin only in the apical adherens junctions and weakly in the basal junction of the ectoderm, but not in the pharyngeal ectoderm, in line with a recent report (Salinas-Saavedra et al., 2018). This could indicate that the basal junctions in the ectoderm and all endodermal junctions are qualitatively different. However, apical adherens junctions in the ectoderm and endoderm have a very similar structure on the ultrastuctural level (Fig. S4). As we observe co-localized actin fibers at these junctions, we assume that either another protein replaces β-catenin or β-catenin is simply not detected at these junctions. Indeed, we note that the antibody also fails to stain nuclear β-catenin after early cleavage stages. Therefore, as a cautionary note, we cannot fully rule out that the failure to stain β-catenin in the pharynx and the endoderm is due to technical problems.

**Fig. 12.**
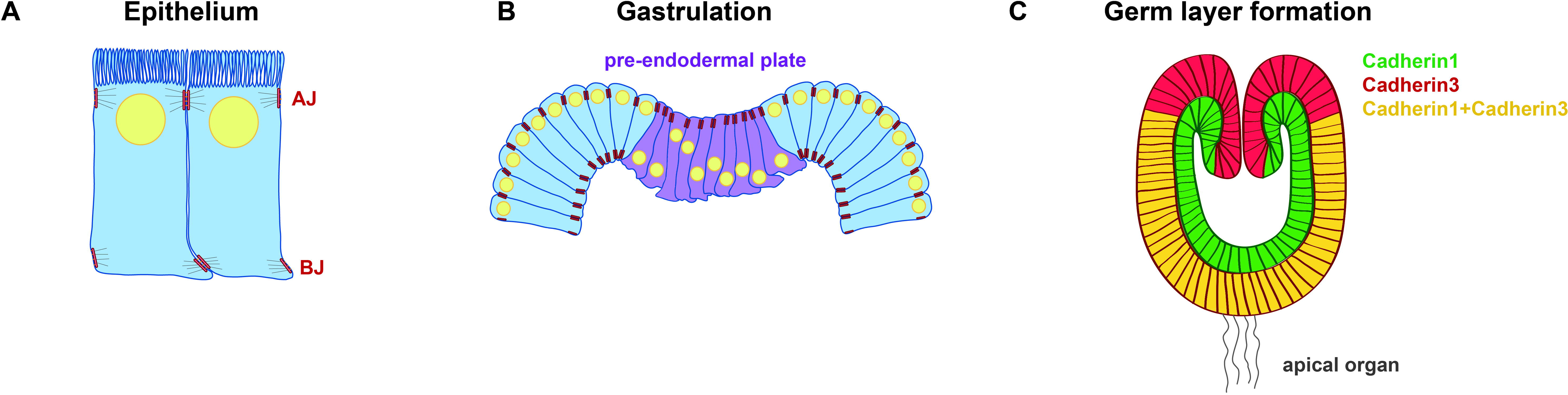
Cadherin localization during early development of *Nematostella*. (A) Schematic depiction of the apical and basal adherens junctions in both epithelial cell layers. (B) The onset of gastrulation is characterized by a downregulation of Cdh3 in the basal junctions, accompanied by apical constriction, migration of nuclei to basal positions, formation of filopodia. (C) Overlapping and specific expression domains of Cdh1 and Cdh3 in a planula larva.

### Cadherin and formation of epithelia

The establishment of the adherens junctions is crucial for normal development of the embryo. Knockdown of *cdh3* in normal embryos does not lead to a dissociation of the embryonic tissue, suggesting that maternally expressed cadherin protein, localized in cell junctions might have a slow turnover and be sufficient for the early stages of development. This is consistent with knockdown of E-Cadherin in mouse embryos (Capaldo and Macara, 2007). However, proper formation of epithelial layers is disrupted in embryonic aggregates upon knockdown of *cdh3*. Notably, while knockdown of the endodermal *cdh1* does not disrupt gastrulation itself, the endoderm does not develop endodermal structures, such as mesenteries. Thus, proper development of the inner germ layer is dependent on the expression of Cdh1.

### The role of cadherins in germ layer formation

The role of cadherins in the formation of germ layers in a diploblast animal is of particular interest, as we might learn about the evolution and potential homology of germ layers. We find that the formation of the inner layer is accompanied by a step-wise cadherin switch. At the blastula stage, Cdh3 forms apical and basal adherens junctions. The onset of gastrulation is characterized by a change of shape of the endodermal cells, which adopt a partial EMT phenotype: They apically constrict, lose the Cdh3-positive basal junctions, the nucleus migrates basally, the cells develop filopodia and become rather motile (Fig. 12B). We propose that the changes in the adhesion properties of the endodermal cells are crucial for the morphogenetic behavior and further differentiation. In a second step, after completion of invagination, Cdh3 also disappears from the apical junctions in the endoderm and is replaced by Cdh1 both at apical and basal junctions of the endoderm. Thus, we observe a cadherin switch in *Nematostella* that is analogous to the cadherin switch in vertebrates and insects. As *cdh1* and *cdh3*, like E- and N-cadherins in vertebrates and insects are lineage-specific duplications, we conclude that the cadherin switch evolved convergently in these animals. However, while Cdh3 is not expressed in the endoderm after the gastrula stage, Cdh1 shows partially overlapping expression with Cdh3 in the ectoderm. Cdh1 seems to form a decreasing gradient from aboral to oral, but the significance of this gradient is unclear at this point. Notably, the oral region and tentacles are completely devoid of Cdh1 expression. We conclude that like in bilaterians (Basilicata et al., 2016; Giger and David, 2017; Huang et al., 2016; Nakagawa and Takeichi, 1998; Pla et al., 2001; Schäfer et al., 2014; Shoval et al., 2007), different combinations and concentrations of Cdh1 and Cdh3 convey different tissue properties and identities in different regions of the developing embryo. Thus, the combinatorial and differential use of cadherins is an recurring feature of metazoans (Fig. 12C, although the molecules have evolved independently.

### Homology of germ layers

Our study has established that cadherins play an important role in the formation and differentiation of the germ layers in a diploblastic animal. This revives the question to which germ layers in Bilateria these two cell layers are homologous. Traditionally, they have been homologized with endoderm and ectoderm, with the mesoderm missing. The identification of a number of mesodermal transcription factors in cnidarians and their expression in the endoderm led to the notion of an inner “mesendoderm” (Fritzenwanker et al., 2004; Kumburegama et al., 2011; Martindale, 2004; Salinas-Saavedra et al., 2018; Scholz and Technau, 2003). However, a recent analysis of many endodermal and mesodermal marker genes has suggested that segregation has already taken place in the *Nematostella* polyp. In fact, the inner layer rather corresponds to mesoderm, while all endodermal functions reside in the ectodermally derived extensions of the pharynx, the septal filaments (Hashimshony, 2017; Steinmetz et al., 2017). In the light of these new findings, it is interesting to note that cdh1 is specific to the cell layer, which would correspond to the mesoderm of bilaterians. Notably, this cell layer also expresses *snail* transcription factors, which regulate the downregulation of *E-cadherin* in vertebrates and insects in the ingressing mesoderm (Fritzenwanker et al., 2004; Martindale, 2004). In line with this, *snail* genes appear to play a role in regulating invagination and partial EMT in *Nematostella* (Salinas-Saavedra et al., 2018). It will be of interest to investigate how the cadherins are regulated by Snail in *Nematostella*.

### Conclusion

This first analysis of the expression and function of classical cadherins in a diploblast shows that these molecules play a conserved role in cell adhesion, tissue morphogenesis and germ layer specification during embryogenesis. The invaginating cells show partial EMT, accompanied by a cadherin switch. The evolutionarily recurring mechanism of a cadherin switch suggests that the evolution of germ layer formation and tissue morphogenesis was facilitated by the differential expression of cadherins.

## Material and Methods

### Animals and embryo culturing

Animals were kept in artificial seawater at 18°C the dark. Spawning was induced by temperature shift to 24°C and light exposure over 10 hours (Fritzenwanker and Technau, 2002). In vitro fertilized embryos were collected and kept at 21 °C as described (Fritzenwanker and Technau, 2002; Genikhovich and Technau, 2009c).

### Identification of Cdh1 and Cdh3 protein sequences

To retrieve the coding sequences of *cdh1* and *cdh3* genes the 1-3Kb overlapping CDS pieces of *cdh1* and *cdh3* were amplified from the cDNA of the mixed embryonic stages, cloned using pJet1.2/blunt vector system (ThermoFisher Scientific) and sequenced. The full-length sequence of Cadherin1 and Cadherin3 have been deposited at Genbank. Assembled *cdh1* and *cdh3* protein coding sequences were translated into the proteins using Expasy translation tool (Artimo et al., 2012). Cadherin protein domain annotation was performed using SMART protein domain annotation resource (Letunic and Bork, 2018).

### Morpholino injection

*cdh1* and *cdh3* knockdowns were performed by independent zygote injections of two nonoverlapping translation blocking morpholinos (Gene Tools, LLC):

cdh1MO1 - 5’ CCGGCCAGCACTCATTTTGTGGCTA 3’;
cdh1MO2 - 5’ ACCCGTGAGTTTAAAAACCCATAGC 3’;
cdh3MO1 5’ ACGAGTTGCGGTGAACGAAAATAAC 3’;
cdh3MO2 5’ TAGCAGAACCGTCCAGTCCCATATC 3’ at concentration 500 μM.
Standard morpholino injection at 500 μM was used as a control.
SdtMO - 5’ CCTCTTACCTCAGTTACAATTTATA’

Non-overlapping morpholinos for *cdh1* and *cdh3* knockdown gave similar effects. Injection equipment used: FemtoJet (Eppendorf), CellTram Vario (Eppendorf), micromanipulator (Narishige), needles were pulled from the glass capillaries type GB 100TF-10 (Science Products GmbH) with a micropipette puller (Sutter Instrument CO., Model P-97). We used holding capillaries from Eppendorf for the injection.

#### Short hairpin RNA (shRNA) knockdown

Target for shRNA *cdh1* knockdown was identified and *cdh1* shRNA design and synthesis were performed as described (He et al., 2018). The following primers were used for *cdh1* shRNA synthesis:

*cdh1* shRNA F: TAATACGACTCACTATAGAAGCGCGCTCAGGTAAATGTTTCAAGAGA
*cdh1* shRNA R: AAGAAGCACGTTCGGGT AAAT GTTCTCTTGAAACATTT ACCTGAGCGC

Purified shRNA injected into zygotes at concentration of 500 ng/μl. As a positive control shRNA against mOrange was injected at 500 ng/μl. Uninjected embryos from the same batch were used as a negative control. After the injection embryos were raised at 21 °C.

### Antibody generation

For anti-Cdh1 antibody generation we expressed in *E.coli* and column-based affinity purified recombinant proteins for immunization. Protein domains cdh1 domain1 (extracellular) and cdh1 domain3 (intracellular) were used for polyclonal antibody production in rats and rabbits respectively (see recombinant sequences in the SI section). Both antibodies resulted in the same staining pattern.

For visualization of Cdh3, monoclonal antibodies were produced in mice. The following peptides were used for immunization: SSSDRNRPPV; DEKDPPQFSQ. Epitopes are located in the extracellular part of Cdh3 in the third and seventh cadherin repeats (CA) respectively. Both antibody clones resulted in the same staining patterns.

### Antibody and Phalloidin staining

For Cdh1 antibody staining embryos were fixed for 1 h at 4 °C with Lavdovsky’s fixative (3,7% formaldehyde (FA)/50% ethanol/4% acetic acid). For cdh3 antibody and phalloidin staining embryos were fixed for 1 h with 3,7% FA in PBS at 4 °C. Primary polyps were relaxed prior the fixation by putting them in 0,1 M MgCl_2_ in *Nematostella* medium for 10 min. After the fixation embryos were incubated on ice with the ice cold acetone (chilled at - 20°C) for 7 min followed by 5 washes with PBSTx 0,2% (PBS with 0,2% of TritonX-100).

After that embryos were incubated in blocking solution (20% sheep serum, 1% Bovine Serum Albumin (BSA) in PBSTx 0,2%) for 2 h at room temperature (RT). Primary mouse anti-Cdh3 antibodies (1:1000) and rat/rabbit anti-Cdh1 antibodies (1:500) were diluted blocking solution and incubated with the embryos overnight at 4 °C. Followed by 10 X 10 min washing steps in PBSTx 0,2% at RT. After incubation in blocking solution for 2 h at RT, embryos were placed in a secondary antibody solution: goat anti-mouse Alexa Fluor 568 antibodies (1:1000), goat anti-rat Alexa Fluor 488 antibodies (1:1000) and DAPI (1:1000) overnight at 4 °C. When fixed with FA, phalloidin Alexa Fluor 488 (1:30) (Thermo Fisher Scientific) was added to the secondary antibody solution, since phalloidin staining is not compatible with the Lavdovsky’s fixation. Followed by 10 X 10 min washing steps in PBSTx 0,2% at RT, embryos were infiltrated with Vectashield antifade mounting medium (Vector laboratories) at 4 °C overnight. Imaging was performed with Leica TCS SP5 DM-6000 confocal microscope.

### *In situ* hybridization

*In situ* hybridizations of embryos were conducted as previously described (Genikhovich and Technau, 2009b; Kraus et al., 2016). The following regions of the CDS of cadherins were used to produce the *in situ* hybridization probes: for *cdh1* (7054bp…9126bp); for *cdh3* (2728bp…5091bp). Adult animals and juveniles were relaxed for 20 min in 0,1M MgCl_2_ solution in *Nematostella* medium followed by fixation and *in situ* hybridization as described (Steinmetz et al., 2017). After *in situ* hybridization adult and juvenile pieces were embedded in 10% gelatine in PBS. Gelatine blocks were post-fixed in 3,7 % FA in PBS overnight at 4 °C and sectioned on a vibratome. Embryos and adult and juveniles sections were embedded in 80% glycerol and imaged with the Nikon Eclipse 80i compound microscope equipped with DIC optics and Zeiss AxioCam camera.

### Time-lapse microscopy

Time-lapse imaging was carried out with the use of a Nikon Eclipse 80i compound microscope. Pictures were taken Zeiss AxioCam camera. Time-lapse movies were made with the use of FIJI software (Schindelin et al., 2012).

### Transmission electron microscopy (TEM)

TEM was performed as previously described (Fritzenwanker et al., 2007).

### Image processing

Images were processed and adjusted for brightness and contrast using FIJI software (Schindelin et al., 2012). Focus stacking of ISH images was done using Helicon Focus software (Helicon Soft Ltd, Kharkov, Ukraine). Images were cropped and assembled into the figures as well as schemes were made using Adobe Illustrator CS6 software (Adobe, San Jose, USA).

## Acknowledgements

We thank Eduard Renfer and Sarah Streinzer for cloning the *in situ* probes of *cdh1* and *cdh3* and performing first *in situ* hybridization of these genes. We thank Dr. Robert Zimmermann for performing the statistical analysis of the aggregate sizes. We thank Vienna Biocenter Facilities (VBCF) for support with recombinant cadherin protein generation and purification. We thank the Core Facility for Cell Imaging and Ultrastructure Research of the University of Vienna (CIUS) for assistance. This work was funded by a Austrian Science Fund (FWF) grant to U.T. (P25993).

## Author contributions

E.P, U.T. conceived the study; E.P. and U.T. designed the experiments; E.P. and A.K. performed the experiments; Y.K. and A.K. performed transmission electron microscopy imaging; E.P. and U.T. wrote the paper.

